# *Sarrella massiliensis* gen. nov. sp. nov., a new gut genus isolated from the stool of a healthy donor

**DOI:** 10.64898/2026.11.03.764080

**Authors:** Rayan Semmar, Mamadou Beye, Nicolas Orain, Simon Robinne, Stéphane Alibar, Sofiane Bakour, Jean-Christophe Lagier, Grégory Dubourg, Maryam Tidjani-Alou, Camille Valles

## Abstract

Strain Marseille-QA0792 (=CSUR QA0792, CECT 31199) was isolated from the feces of a healthy person using the culturomics method. The fecal sample originated from leftover material collected for a fecal microbiota transplantation (FMT) procedure, obtained during a routine screening in accordance with the guidelines of the French National Agency for Medicines and Health Products Safety (ANSM). This strain is Gram-stain-negative, rod-shaped, non-motile, and non-spore-forming. It is strictly anaerobic, with optimal growth at 37°C. Strain Marseille-QA0792 presented its highest 16S rRNA identity, 96.2%, with “*Merdibacter massiliensis”* strain Marseille*-*P3254 (not validly published) (accession number LT598590) and ANI similarity of 75.0% with the same species. Among validly published species, strain Marseille-QA0792 showed its highest 16S rRNA gene sequence similarity (91.5%) with *Faecalicoccus acidiformans* strain 37-2a (accession number HQ452864), whereas all computed average nucleotide identity (ANI) values were below 75%. Genome sequencing revealed a genome size of 2.67 Mb and a G+C content of 50.01%. This species is classified within the *Erysipelotrichaceae* family in the Bacillota phylum. Thus, we isolated for the first time a novel genus named *Sarrella massiliensis* gen. nov., sp. nov., strain Marseille-QA0792. This discovery highlights the importance of continuing culturomics efforts to explore the human microbiome.

## Introduction

Culturomics has enabled the discovery of numerous commensal bacteria from various human microbiotas [1]. The gut microbiota is the richest and is associated with numerous human pathologies [2]. Culturing microbes therefore proves fundamental to better understanding its functioning [3]. Indeed, certain bacteria isolated from the human gut have proven to be of major interest for human health. First, there was *Akkermansia muciniphila*, which plays a major role in the intestinal mucosa [4], followed by *Faecalibacterium prausnitzii* and *Christensenella minuta*, key bacteria associated with good human health [5,6]. More recently, *Dysosmobacter welbionis* (initially discovered and cultured under the name “Oscillibacter massiliensis”) [7] was identified as a bacterium present in 70% of the general population, whose fecal abundance is negatively correlated with body mass index, fasting glucose, and glycated hemoglobin in obese subjects with metabolic syndrome [8]. In this context, it is essential to continue culturing gut microbiota bacteria to identify other bacteria of interest for human health. Thus, using new culture conditions in culturomics, we isolated a new bacterial genus belonging to the family *Erysipelotrichaceae* and the phylum Bacillota. This family comprises numerous genera validly published on the List of Prokaryotic names with Standing in Nomenclature (LPSN), with *Erysipelothrix* as the type genus.

Thus, we classify and describe this previously uncultivated strain isolated using the culturomics technique with the taxonogenomics approach, a polyphasic strategy combining phenotypic and phylogenomic characteristics [9].

## Materials and methods

### Strain isolation and ethics statement

Strain Marseille-QA0792 was isolated from a frozen stool sample obtained from a healthy person, corresponding to residual material from a fecal microbiota transplantation (FMT) procedure. This sample had originally been collected for routine screening of gastrointestinal pathogens, in accordance with the recommendations of the French National Agency for Medicines and Health Products Safety (ANSM) [10]. Eligible subjects were aged between 18 and 65 years, with a body mass index below 30. Exclusion criteria comprised recent antibiotic treatment, travel to intertropical regions within the three months before donation, prolonged residence (several years) in intertropical regions, and hospitalization abroad exceeding 24 hours within the previous 12 months. Written informed consent was obtained from the stool donor. Ethical approval for the study was granted by the Ethics Committee of the Assistance Publique-Hôpitaux de Marseille (AP-HM) under reference CSE 24-4. This isolate, together with six other new species identified from the same stool specimen, was recovered as part of a study seeking an alternative to rumen fluid for the enrichment of blood culture bottles. Several enrichment additives, including fecal filtrate and mucin, were tested, and this strain was obtained using an anaerobic blood culture bottle (bioMérieux, Craponne, France) supplemented with mucin (Sigma-Aldrich, St Louis, USA), sterile rumen fluid (0.2 μm-filtered) and defibrinated horse blood (Thermo Scientific, Hampshire, UK). The sample was seeded after dilution on 5% sheep blood-enriched Columbia (COS) agar (bioMérieux, Craponne, France), and the strain Marseille-QA0792 was isolated after picking on 5% sheep blood-enriched Columbia (COS) agar (bioMérieux, Craponne, France). Identification of the previously isolated colony using a SiriusTM MALDI-TOF mass spectrometer (Bruker Daltonics, Bremen, Germany) with the MALDI Biotyper HT software associated with BDAL micro-organisms library (v.13 Revision G) proved inconclusive (score <1.7, below the threshold defined for reliable identification: >2.0) [11].

### Genomic analysis

The strain was sequenced using a hybrid approach combining MiSeq technology (Illumina Inc, San Diego, CA, USA) with the paired-end strategy, and GridION technology (Oxford Nanopore Technologies), the latter being used to fill gaps in the Illumina assembly. Raw sequencing reads were first subjected to quality control using FastQC v0.11.9 [12], followed by trimming and quality improvement with Trimmomatic v0.39 [13]. Genome assembly was then performed using Unicycler v0.4.8 [14]. Scaffolds shorter than 800 bp were discarded to reduce the risk of contamination [15], and assembly quality was evaluated with CheckM [16]. The 16S rRNA gene sequence was identified and extracted using Barrnap [17]. Genomic and biochemical characterization of the strain will be performed using Protologger v2 [18,19] (https://doi.org/10.5281/zenodo.21694383). To assess whether the strain represents a new species, several taxonomic markers were used, including 16S rRNA gene sequence similarity, with a species delineation threshold of 98.65% [20] and 94.5% for new genera [21], Average Nucleotide Identity (ANI) with a species delineation threshold set at 95% [22], G+C percent difference > 1% [23], and Percentage Of Conserved Protein (POCP) with a threshold set at 50% for genus delineation [24].

### Phylogenetic trees based on the 16S rRNA gene and the genome

To classify the strain, phylogenetic analyses of the genomes and 16S rRNA genes were performed in IQ-TREE2 [25] using the Maximum Likelihood method and the General Time Reversible (GTR) model (Nei & Kumar, 2000). Node robustness was evaluated through 1000 bootstrap replicates, and the proportion of replicate trees recovering the same taxa grouping was shown alongside each branch [27]. Sequence alignments were performed with MAFFT [28] for 16S rRNA genes and with Mugsy version 1.2.3 [29] for genomes. The resulting phylogenetic trees were visualized with iTOL [30], and accession numbers for the 16S rRNA sequences of type strains included in the tree construction were obtained from LPSN.

### Phenotypic and biochemical characterization

Optimal growth conditions for the strain were determined by testing several temperatures (room temperature (RT), 28, 37, 45 and 56°C) and atmospheres (aerobic, microaerophilic and anaerobic) on COS agar plates (bioMérieux). Anaerobic and microaerophilic atmospheres were generated using GasPak™ EZ anaerobic and microaerophilic generators (Becton Dickinson, Franklin Lakes, New Jersey, USA), respectively. Tolerance to pH and salinity was assessed by testing growth at various pH values (5, 5.5, 6, 6.5, 7, 7.5, 8, 8.5) and NaCl concentrations (7.5%, 10%, 15% and 20%). Phenotypic characterization further included Gram staining, spore-forming ability and motility, as well as oxidase (bioMérieux) and catalase (Becton Dickinson, Franklin Lakes, NJ, USA) tests [31]. Furthermore, the metabolic profile of the strain was determined using API ZYM, API 50CH and API 20A strips (bioMérieux)

### Fatty acid methyl ester (FAME) profiling using Gas Chromatography/Mass Spectrometry (GC/MS)

Cellular FAME profiling was performed using GC/MS. Two samples were prepared with approximately 25 mg of bacterial biomass per tube, harvested from several culture plates. Fatty acid methyl esters were prepared as described [32]. GC/MS analyses were carried out as described before [33]. Briefly, fatty acid methyl esters were separated using an Elite 5-MS column and monitored by mass spectrometry (Clarus 690 – SQ8T, Perkin Elmer, Courtaboeuf, France). Spectral database search was performed using MS Search 2.0 operated with the Standard Reference Database 1A (NIST, Gaithersburg, USA) and the FAMEs mass spectral database (Wiley, Chichester, UK).

## Results

### Phenotypic and biochemical analysis

Strain Marseille-QA0792 grew optimally at 37°C under strict anaerobic conditions, and growth was also observed under microaerophilic conditions. This strain exhibited good growth across a pH range of 5.5 to 7.5 but did not tolerate saline environments. It was a Gram-stain-negative, rod-shaped bacterium, non-motile and non-spore-forming, forming white colonies of 1 mm in diameter with regular, convex margins. This strain was catalase and oxidase-negative and able to metabolize glycerol, D-arabinose, L-arabinose, D-ribose, L-xylose, D-adonitol, D-galactose, D-glucose, D-fructose, D-mannose, L-sorbose, L-rhamnose, dulcitol, inositol, D-mannitol, D-sorbitol, methyl-αD-mannopyranoside, methyl-αD-glucopyranoside, N-acetylglucosamine, amygdalin, arbutin, esculin ferric citrate, salicin, D-cellobiose, D-maltose, D-lactose, D-melibiose, D-saccharose, D-trehalose, inulin, D-melezitose, D-raffinose, glycogen, xylitol, D-tagatose, D-fucose, L-fucose, D-arabitol, L-arabitol, potassium gluconate, and 5-ketogluconate. Using the API ZYM strip, enzymatic activities were detected for acid phosphatase, naphthol-AS-BI-phosphohydrolase, and α-glucosidase. Using the API 20NE strip, arginine dihydrolase, urease, and esculin hydrolase activities were also detected, and glucose fermentation was positive. The major fatty acids were Hexadecanoic acid (26.9%), 9-Octadecenoic acid (22.0%), and Octadecanoic acid (18.0%). Several molecules with shorter (C10) or longer carbon chains (C20) were detected. No branched-chain structures were identified.

### Strain identification and taxonomic analysis

MALDI-TOF MS analysis of strain Marseille-QA0792 yielded qualitative spectra but did not allow species-level identification (Figure 1). Strain Marseille-QA0792 ^T^ showed its highest 16S rRNA sequence identity (96.2%) with “;*Merdibacter massiliensis*” strain Marseille-P3254 (not validly published) (accession number LT598590), a result confirmed by the 16S rRNA phylogenetic tree (Figure 2), along with an ANI similarity of 75.0% to the same species, further supported by the genome-based phylogenetic tree (Figure 3). Based on the Protologger analysis (validly published species only), the 16S rRNA sequence showed the highest similarity to *Faecalicoccus acidiformans* strain 37-2a ^T^ (91.5%; accession number HQ452864). All calculated ANI and POCP values remained below the species (75%) (Figure 4) and genus (50%) delineation thresholds, respectively. These analyses support the classification of strain Marseille-QA0792 as a new species within a novel genus.

**Figure 1.**
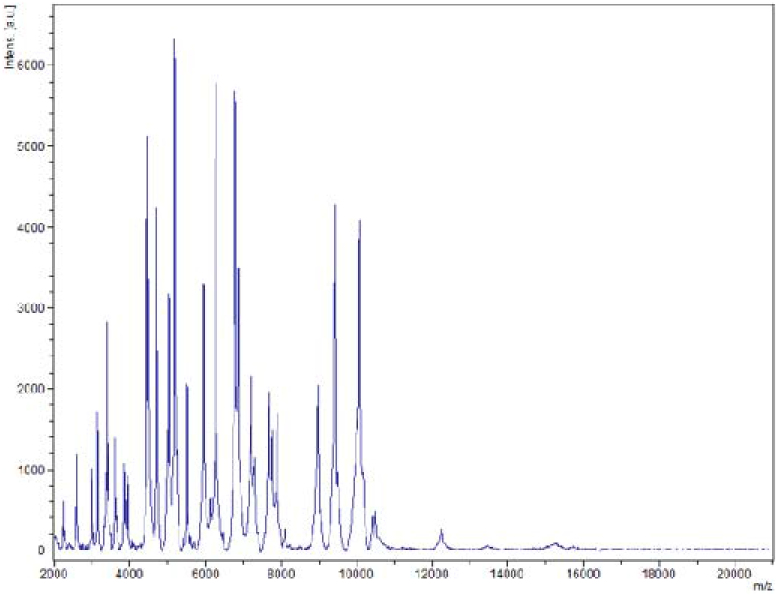
MALDI-TOF spectrum (smoothed and baseline-subtracted) of strain Marseille-QA0792, generated using FlexAnalysis v3.4 (Bruker) software.

**Figure 2.**
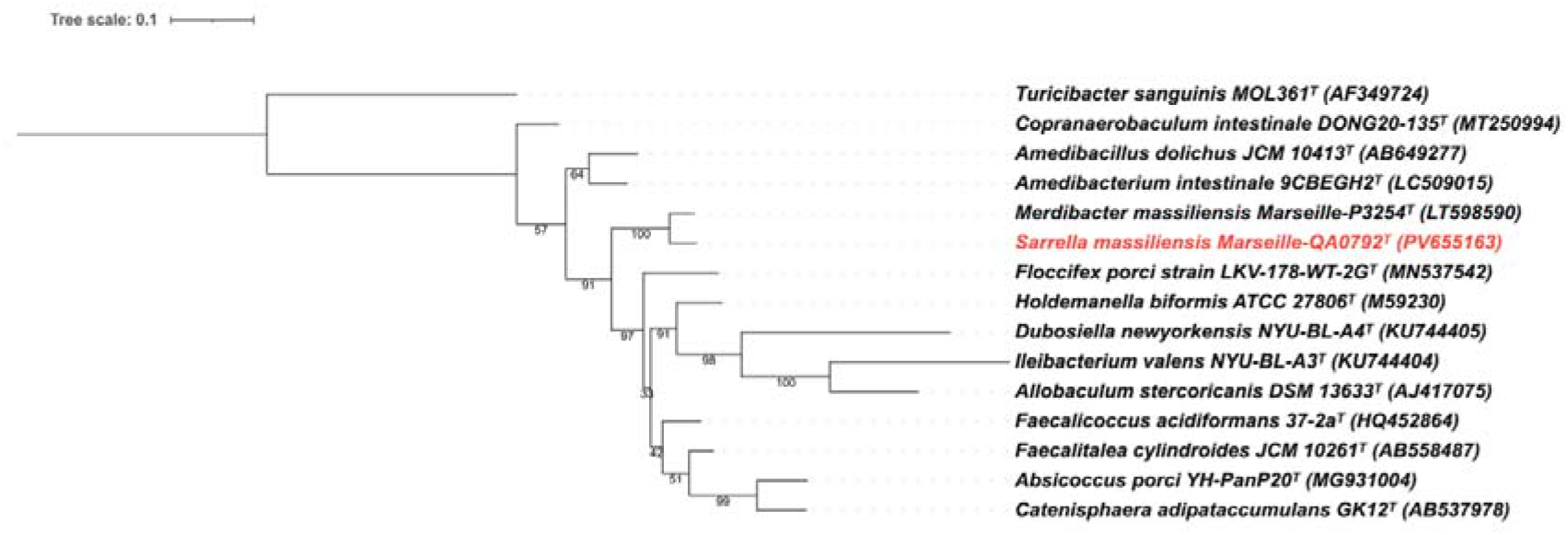
Phylogenetic tree based on the 16S rRNA gene, constructed using the Maximum Likelihood method with the General Time Reversible model and 1000 bootstrap replicates. *Turicibacter sanguinis* (family *Turicibacteraceae*) was used as the outgroup. Collection numbers and accession numbers are indicated after each species name.

**Figure 3.**
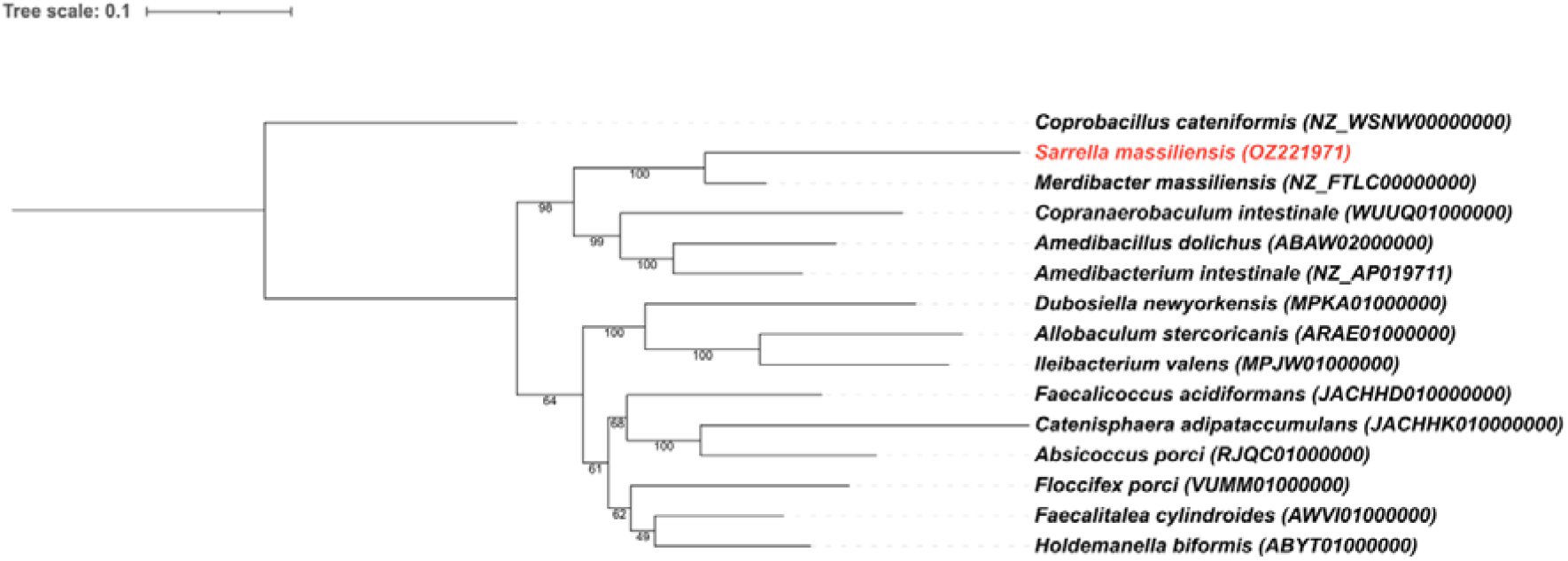
Genomic tree constructed using the Maximum Likelihood method with 1000 bootstrap replicates. *Coprobacillus cateniformis* (family *Coprobacillaceae*) was used as the outgroup. Collection numbers and accession numbers are indicated after each species name.

**Figure 4.**
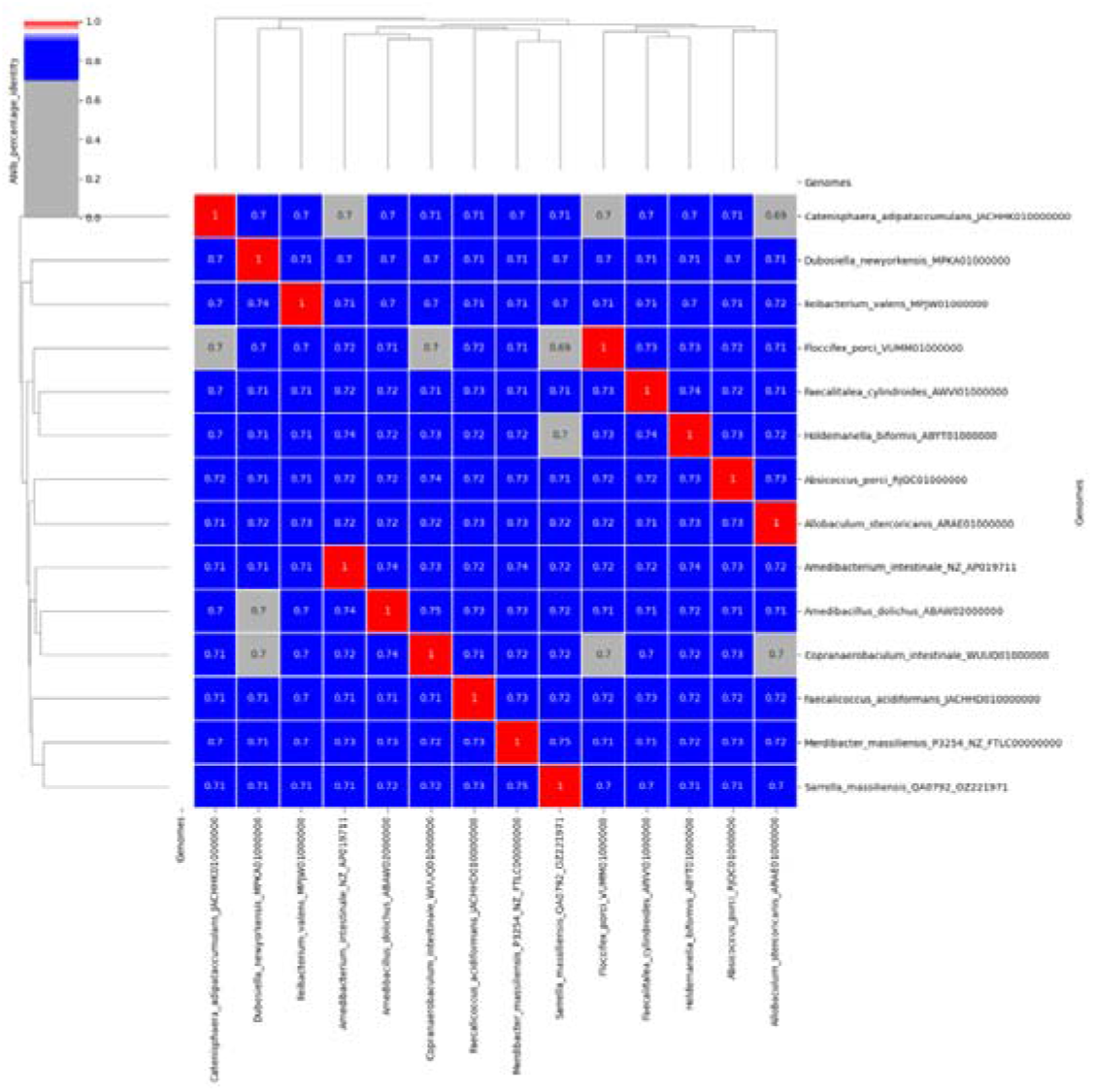
Heatmap showing the ANI similarity percentages among species related to *Sarrella massiliensis*.

### Genomic annotation

The genome of strain Marseille-QA0792 comprised a single contig of 2.67 Mbp with a G+C content of 50.01%. Genome quality assessment by CheckM revealed high completeness (99.06%) and low contamination (1.89%). A total of 2,471 protein-coding genes were identified by Protologger analysis, including 120 transporters, 16 secretion genes, 520 unique enzymes, and 145 CAZymes. The annotation of our genome using the RAST software (https://rast.nmpdr.org/<u>, last accessed July, 2026</u>) [34] revealed the presence of 64 RNAs, including 56 tRNAs, four 23S rRNAs, and four 16S rRNAs. Biochemical prediction performed via Protologger, based on KEGG (Kyoto Encyclopedia of Genes and Genomes) annotations, highlighted several pathways encoded within the genome. Propionate production was predicted through two independent pathways: from propanoyl-CoA (EC:2.8.3.1) and via a second pathway (GMM:MF0094), while ammonia was predicted to be converted into L-glutamate through an L-glutamine intermediate (EC:6.3.1.2, 1.4.1.-). Fermentation-associated functions were also detected, including lactate synthesis and the conversion of acetyl-CoA into acetate (GMM:MF0092; GMM:MF0086). The genome further encoded biosynthetic capacities for a range of amino acids: proline, cysteine, ornithine, valine, isoleucine, leucine, phenylalanine, tyrosine and arginine, as well as for coenzyme A and several cell envelope building blocks, including UDP-glucose, UDP-GlcNAc, UDP-MurNAc, UDP-ManNAcA and UDP-GalNAcA. Both glycolysis and fatty acid biosynthesis pathways appeared complete, supporting their likely functionality.

## Conclusion

We report the isolation of a previously uncultured species from the feces of a healthy individual. Strain Marseille-QA0792 was characterized using the taxonogenomics approach and formally named *Sarrella massiliensis* gen. nov., sp. nov. This finding underscores the importance of optimizing culturomics approaches to further reduce the fraction of bacterial diversity that remains uncultured.

### Description of *Sarrella* gen. nov

*Sarrella* gen. nov (Sar.rel’la. N.L. fem. dim. n. *Sarrella*, in honor of the senegalese microbiologist Salematou Sarr). Strain Marseille-QA0792 exhibits its highest overall 16S rRNA sequence identity (96.2%) with “*Merdibacter massiliensis*” strain Marseille-P3254 (not validly published; accession number LT598590) and an ANI similarity of 75.0% with the same species. When considering only validly published species, Protologger analysis identifies *Faecalicoccus acidiformans* strain 37-2a as the closest relative, with a 16S rRNA sequence identity of 91.5% (accession number HQ452864). Notably, all calculated ANI and POCP values remain below 75% and 50%, respectively. The type strain of *Sarrella* gen. nov. is Marseille-QA0792 (CSUR QA0792=CECT 31199), which was isolated from the feces of a healthy person. The 16S rRNA and genome are deposited on the NCBI platform under accession numbers PV655163.1 and OZ221971.1, respectively

### Description of *Sarrella massiliensis* sp. nov

*Sarrella massiliensis* sp. nov (mas.si.li.en’sis. L. fem. adj. *massiliensis,* of Massilia, the Latin name of Marseille). Optimal growth is observed at 37°C in a strict anaerobic atmosphere on 5% sheep blood-enriched Columbia agar. This species is a non-motile, non-spore-forming, Gram-stain-negative, and rod-shaped bacterium. This species forms white colonies, 1 mm in diameter, with regular, convex margins and grows well at pH levels ranging from 5.5 to 7.5 but shows no tolerance for saline environments. It can metabolize glycerol, D-arabinose, L-arabinose, D-ribose, L-xylose, D-adonitol, D-galactose, D-glucose, D-fructose, D-mannose, L-sorbose, L-rhamnose, dulcitol, inositol, D-mannitol, D-sorbitol, methyl-αD-mannopyranoside, methyl-αD-glucopyranoside, N-acetylglucosamine, amygdalin, arbutin, esculin ferric citrate, salicin, D-cellobiose, D-maltose, D-lactose, D-melibiose, D-saccharose, D-trehalose, inulin, D-melezitose, D-raffinose, glycogen, xylitol, D-tagatose, D-fucose, L-fucose, D-arabitol, L-arabitol, potassium gluconate, and 5-ketogluconate. The following enzymatic activities are detected: acid phosphatase, naphthol-AS-BI-phosphohydrolase, α-glucosidase, arginine dihydrolase, urease, and esculin hydrolase. Glucose fermentation is also positive. The most abundant fatty acid is Hexadecanoic acid (26.9%). The KEGG analysis identifies putative metabolic pathways for this strain, such as the production of propionate and L-glutamate. This strain can utilize cellobiose, sucrose, and starch as carbon sources. The genome size is 2.67Mbp with a G+C content of 50.01%. The type strain of *Sarrella massiliensis* gen. nov., sp. nov. is Marseille-QA0792 (CSUR QA0792=CECT 31199), which was isolated from the feces of a healthy person. The 16S rRNA and genome are deposited on the NCBI platform under accession numbers PV655163.1 and OZ221971.1, respectively.

### Strain deposit

Strain Marseille-QA0792 was deposited in the Collection des Souches de l’Unité des Rickettsies (CSUR) under deposition numbers CSUR QA0792 and deposited in the Spanish Type Culture Collection (CECT) under number CECT 31199.

## Data availability

The 16S rRNA gene and genome sequences are available in the NCBI database under accession numbers PV655163.1 and OZ221971.1, respectively. Protologger data for *Sarrella massiliensis* strain Marseille-QA0792 were deposited on Zenodo (https://doi.org/10.5281/zenodo.21694383).

## Authors’ contributions

**Conceptualization:** R.S, C.V. **Formal analysis:** M.B **Investigation:** R.S, N.O. **Methodology:** M.T.A. **Resources:** S.A, S.B. **Supervision:** C.V **Validation:** M.T.A, G.D, C.V, JC.L. **Visualization:** R.S, S.R, M.B, S.B, N.O. **Writing-Original draft:** R.S, M.B, N.O. **Writing: Review & Editing:** M.T.A., C.V, G.D, JC.L

## Acknowledgements

We thank the CSUR team for strain preservation and Dr Ousmane NDOUR for the microscope assistance. We thank Dr. Aurélia Caputo for the genome and 16S rRNA sequence deposit at the European Bioinformatics Institute (EBI).

## Ethical approval and consent to participate

The stool sample was obtained as part of a fecal microbiota transplantation procedure, in accordance with the recommendations of the French National Agency for the Safety of Medicines (ANSM) (https://ansm.sante.fr/uploads/2022/09/19/microbiote-fecale-rapport-nov-2016.pdf). Investigation of bacterial species within this stool specimen was approved by the Ethics Committee of the Assistance Publique-Hôpitaux de Marseille, under reference number CSE 24-4.

## Availability of data

All data generated or analyzed during this study are included in this published article.

## Declaration of AI-assisted

The authors acknowledge the use of generative artificial intelligence tools for English language editing and manuscript refinement.

---

**Table 1.** Cellular fatty acid composition (%) of strain Marseille-QA0792.

| **Fatty acids** | **Name** | **Mean relative %□** |
|---|---|---|
| 16:0 | Hexadecanoic acid | 26.9 ± 0.7 |
| 18:1n9 | 9-Octadecenoic acid | 22.0 ± 1.3 |
| 18:0 | Octadecanoic acid | 18.0 ± 0.4 |
| 18:2n6 | 9,12-Octadecadienoic acid | 9.8 ± 1.0 |
| 14:0 | Tetradecanoic acid | 9.5 ± 0.2 |
| 18:1n7 | 11-Octadecenoic acid | 8.0 ± 0.3 |
| 16:1n7 | 9-Hexadecenoic acid | 3.3 ± 0.2 |
| 20:1n9 | 11-Eicosenoic acid | TR |
| 12:0 | Dodecanoic acid | TR |
| 15:0 | Pentadecanoic acid | TR |
| 20:4n6 | 5,8,11,14-Eicosatetraenoic acid | TR |
| 14:1n | Tetradecenoic acid | TR |
| 20:1n | Eicosenoic acid | TR |
| 20:0 | Eicosanoic acid | TR |
| 10:0 | Decanoic acid | TR |
| 16:1n9 | 7-Hexadecenoic acid | TR |

*a Mean peak area percentage; TR = trace amounts < 1 %*

